# 53BP1 condensates function as bioreactors for NHEJ directed DNA repair and insulators to determine pathway choice

**DOI:** 10.64898/2025.12.19.695596

**Authors:** Mikael V. Garabedian, Tianpeng Zhang, Arindam Datta, Wentao Wang, Maxwell Wadley, Roger A. Greenberg, Matthew C. Good

**Affiliations:** Department of Cell and Developmental Biology, Perelman School of Medicine, University of Pennsylvania, Philadelphia, PA 19104-6160, USA; Department of Cancer Biology, Penn Center for Genome Integrity, Basser Center for BRCA, Perelman School of Medicine, University of Pennsylvania, Philadelphia, PA 19104-6160, USA; Department of Bioengineering, University of Pennsylvania, Philadelphia, PA 19104-6160, USA

## Abstract

Genomic integrity requires efficient resolution of DNA damage. Non-homologous end joining (NHEJ) is the primary mechanism of DNA double strand break (DSB) repair in mammalian cells and is mediated by 53BP1, a tumor suppressor involved in preventing DSB end-resection and homologous recombination. NHEJ repair foci form following DSB formation, however how mesoscale assembly occurs and whether 53BP1 is the driver of this process are unknown. Also, despite knowledge of the identify of key pathway molecules, the specific functions of mesoscale repair condensates in DNA repair and pathway selectivity are unknown. To address these gaps, we determined the minimal domain of 53BP1 sufficient for phase separation in vitro and identified the key residues that governing its condensation. Utilizing a separation-of-function mutant, we demonstrate that 53BP1and its protein condensation is the core driver of NHEJ foci formation. 53BP1 condensates function as bioreactor compartments, increasing the effective concentration of substrates near double strand breaks and are essential for efficient DNA damage resolution. Additionally, we show that 53BP1 condensates function as insulators around DSB sites to prevent end-resection and direct repair pathway selectivity. Collectively our work reveals a specialized compartment for DNA repair through the spatial clustering of 53BP1 molecules into repair foci essential to maintain genome integrity.

## MAIN TEXT

Cells are exposed to agents that damage DNA and experience hundreds to thousands of different DNA lesions in each cell cycle^1^. Double stranded breaks (DSBs) and single stranded breaks or nicks that spontaneously convert to DSBs are especially hazardous to genomic integrity^1, 2^. Each DSB must be quickly detected and repaired by the appropriate repair pathway to prevent chromosomal loss, translocations, and other aberrations that lead to genomic instability^1, 3^. Defective DNA repair is strongly associated with mutational burden and cellular transformation in cancers. In humans, non-homologous end joining (NHEJ) and homology directed repair (HDR) are the major DSB repair pathways, with NHEJ accounting for repair of ∼ 80% of DSBs^4^. HDR is mediated by BRCA1 and requires end resection and strand invasion of an intact sister chromatid for repair. While HDR is most active in late S and G2 phases of the cell cycle^5, 6^, NHEJ is active in all cell cycle phases is engaged in competition with HDR factors for access and broken end processing of DSBs^7–10^. NHEJ is promoted by 53BP1 via antagonism of end-resection and BRCA1 localization to DSBs^11, 12^.

NHEJ repair foci harboring 53BP1 form rapidly at DSBs following DNA damage. Initially, 53BP1 recognizes break associated nucleosome modifications, H2AK15-ubiquitylation and H4K20-dimethylation, at chromatin flanking DSBs as well as γH2AX^13–18^. However, whether further assembly of mesoscale structures harboring 53BP1 are required for NHEJ function is unknown and represents a fundamental gap in understanding regarding how DSBs are processed. Micron-scale NHEJ foci behave similar to liquid-like biomolecular condensates, undergoing fission and fusion events and showing dynamic recovery after photobleaching^19–22^. The C-terminus of 53BP1 was recently shown to form droplet-like structures via optogenetic clustering with CRY2 in cells^19, 23^, suggesting that 53BP1 may contain sequences that promote self-assembly and condensation. However, the single molecular scaffold or set of factors that primarily drive NHEJ repair foci condensation following DNA damage are not known.

In the present work, we set out determine the contribution of biomolecular condensation to promoting NHEJ mediated DNA repair. To first determine the role of 53BP1 in NHEJ foci formation, we identified the minimal regions of 53BP1 required for phase separation, a necessary step to create a specific separation-of-function mutant for condensate formation. We isolated the minimal domain, which includes an intrinsically disordered region (IDR) and oligomerization domain. We identified the critical amino acids in the minimal region that governs phase separation, allowing separation-of-function tests in cells. The condensation mutant version forms smaller foci and are defective in resolving DSBs. Condensate deficient 53BP1 mutants interact normally to their ligands but form significantly smaller foci and thereby recruit deficient amounts of NHEJ factors to DSBs. These data reveal that 53BP1 self-assembly is essential for foci formation, highlighting its role as a key scaffold to promote NHEJ. Additionally, we found condensate deficient 53BP1 mutants are dysfunctional in antagonizing the reciprocal HDR pathway. Our observations indicate that 53BP1 condensates function as both a bioreactor to promote NHEJ as well as an insulator to prevent access to the DSB by resection machinery and HDR factors. Collectively, this study offers new paradigms for the function of biomolecular condensates in the fidelity and function of genome integrity pathways.

### 53BP1 oligomerization domain and first C-terminal IDR drive phase separation

53BP1 is an ∼2000 aa protein with extensive disordered or low complexity sequences. Its C-terminus is predicted to harbor two intrinsically disordered regions (IDRs), flanked by folded domains involved in binding and localization to sites of DNA breaks and protein-protein interactions (**Fig. 1a**). Previous studies identified the C-terminus of 53BP1 as potential starting point for biochemical characterization^19^. To identify the minimal region in the C-terminus that governs phase separation, we purified the 53BP1 C-terminus (1203-1972 aa) as well as subregions, including one that contains the oligomerization domain (OD) and first C-terminal IDR (OD-IDR1; 1226-1479 aa) and one that contains IDR1 alone (1273-1479 aa). We compared their ability to form droplets in vitro under physiological conditions (**Fig. 1b, c, d**). To our surprise, the OD-IDR1 polypeptide formed droplets in vitro at similar concentrations to the much larger C-terminus, suggesting a similar saturation concentration (C_sat_). In contrast, the IDR1 alone did not form droplets in vitro (**Fig. 1d**). These results reveal that IDR2 and other folded domain in the C-terminus are dispensable and identify OD-IDR1 as the minimal sequence responsible for driving condensate formation. Notably, this domain architecture, utilizing an oligomerization domain linked to an IDR would enforce valency to drive protein phase separation and is similar to other proteins found to be essential for condensate formation^24^.

**Fig 1.**
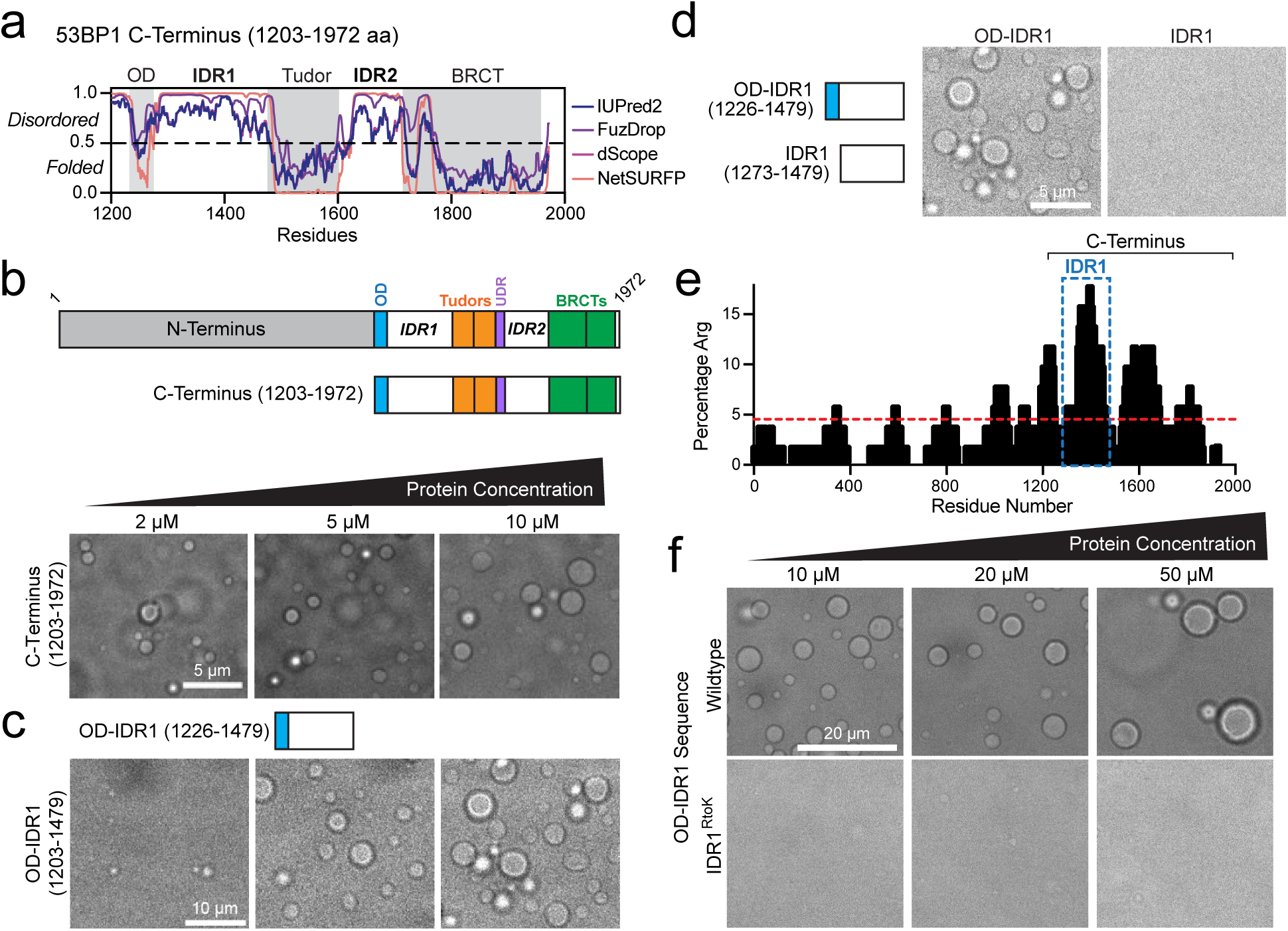
Oligomerization Domain and C-terminal IDR1 comprise minimal region for 53BP1 condensation. **a**, Polypeptide disorder predictions for 53BP1 C-terminus (1203-1972) by multiple algorithms as indicated. Values above 0.5 indicate disorder. Folded domains indicated by values below 0.5 and grey background. **b,** Top: Domain map of full length 53BP1 and the C-terminus, which was purified. Bottom: Purified C-terminus robustly forms droplets in vitro at the indicated concentrations. **c,** Purified polypeptide of the oligomerization domain and IDR immediately downstream (OD-IDR1; 1226-1479) condenses in vitro at the same concentrations as the C-terminus (b). **d,** The oligomerization domain (OD) is required for condensation as IDR1 alone (1273-1479) is not sufficient for condensate formation in vitro. **e,** Sliding window analysis of 53BP1 shows enrichment of arginine (Arg) residues in the 53BP1 C-terminus and most significantly in the C-terminal IDR1 sequence (blue box). Red dotted line indicates typical percentage of Arg in a vertabrate protein sequence. **f,** Converting arginines (R) to lysines (K) in IDR1 disrupts condensation in vitro.

IDR-mediated protein condensation is generally driven by multivalent interactions between amino acids or sequence motifs^25–29^. Specific residue types, such as aromatic or arginine residues have been identified as critical to the phase separation of disordered sequences found in FUS, LAF-1 and hnRPA1^30–32^. To determine whether any portion of 53BP1 is enriched in residues that may participate in multivalent interactions, we analyzed residue enrichment in the 53BP1 sequence. We found an elevated density of arginine (Arg, R) in the C-terminal end of 53BP1 with the largest peak overlapping with IDR1 (**Fig. 1e**). The percentage of Arg in a typical vertebrate protein sequence is 4-5%^33^ (**Fig. 1e**, dotted line in red), whereas the Arg content in IDR1 (blue box) is more than triple this figure. To directly test their role in self-assembly, we purified polypeptides of OD-IDR1 with its wildtype sequence or arginine-to-lysine (R-to-K) mutations in IDR1 (notably, the mutant preserves charge in the IDR sequence and avoids Arg residues within the OD). OD-IDR1 condensed robustly, whereas the R-to-K mutant failed to condense in vitro at any concentration tested (**Fig. 1f, Extended Data Fig. 1a**). This result demonstrates Arg is a key determinant of self-assembly of IDR1. These observations also demonstrate that oligomerization via the OD alone is insufficient for condensation when IDR1 interactions are perturbed. Together, our results reveal the minimal region of 53BP1 sufficient for condensation in vitro and identify key residues for self-assembly which we use moving forward to interrogate the contribution of 53BP1 to repair condensate formation in cells.

### 53BP1 IDR1 mutants disrupt NHEJ foci assembly

Whether 53BP1 is the primary driver of NHEJ foci formation and whether condensate formation contributes to DNA repair function is not known. First, to determine the extent to which 53BP1 drives repair foci formation in in cells, we deleted the native copies in U2OS cells and re-integrated coding sequences to express GFP-tagged full-length 53BP1 and a version containing IDR1 R-to-K mutations (IDR1^RtoK^) as stable cell lines (**Fig. 2a and Extended Data Fig. 2**). Cells were synchronized in G1 phase (when NHEJ is most active), via CDK4/6 inhibition^34^, released, and DNA damage was induced with 2 Gy ionizing radiation (IR). We observed that both wildtype and mutant 53BP1 constructs form nuclear foci that are localized to DSBs marked by γH2AX (**Fig. 2b**). However, foci formed by the IDR1^RtoK^ mutant were markedly reduced in volume compared to the wildtype version (**Fig. 2c**). Further, the mutant displayed diminished partitioning of protein into the condensed phase (**Fig. 2d**), consistent with our in vitro findings. To date the identity of the primary driver of NHEJ foci has been unknown. While 53BP1 is understood to enforce the recruitment of additional machinery, it had not been known whether generation of NHEJ associated foci or their liquid-like properties are specifically dependent on 53BP1 or on one of its many ligands that may also condense (e.g. p53)^35, 36^. Notably, previous studies relied on large deletions and truncations of 53BP1 to characterize its self-assembly, likely leading to pleiotropic effects. By identifying the minimal domain and condensate-defective mutations is it now clear of that 53BP1 is the primary driver of NHEJ foci formation.

**Fig 2.**
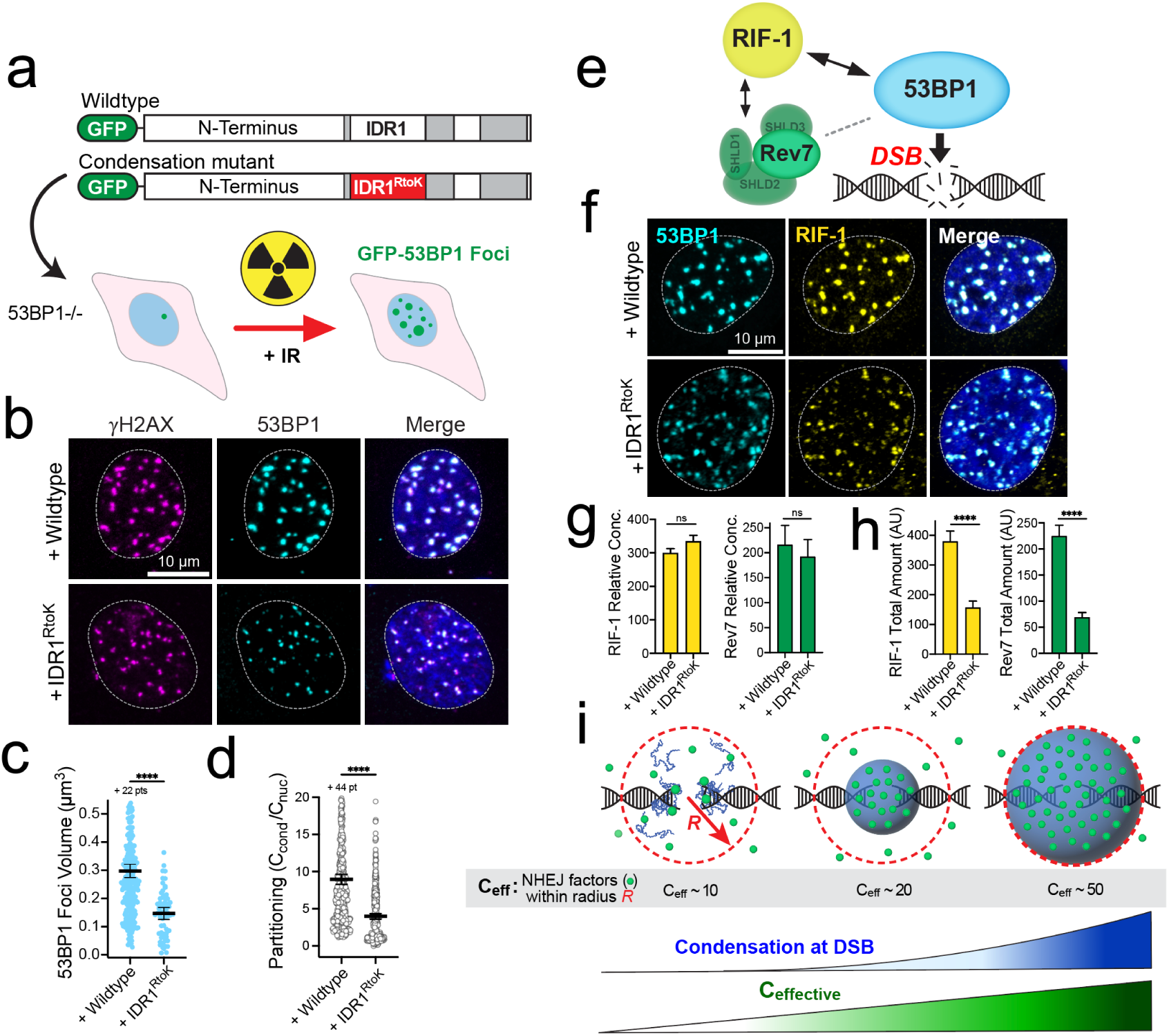
C-terminal IDR1 R-to-K mutations significantly diminish 53BP1 foci volumes in cells. **a**, Experimental schematic. Full length constructs of GFP-53BP1, wildtype or with 13 R-to-K mutations in the C-terminal IDR1, were tranduced into 53BP1−/− U2OS cells. Cells were exposed to 2 Gy IR and 53BP1 foci were evaluated. **b,** Representative immunofluorescence (IF) images of nuclei expressing 53BP1 constructs co-stained for DSBs by gH2AX. **c,** Comparison of 53BP1 foci volumes, wildtype vs IDR1 mutant. **d,** Partitioning of wildtype and IDR1 mutant 53BP1 in foci (condensate) over nucleoplasm. **e,** Diagram of 53BP1-ligand interactions at DSBs. **f,** Representative IF images of nuclei expressing 53BP1 constructs co-stained for 53BP1 binding partner, RIF-1. **g,** measurements of relative concentrations (mean fluorescence per focus) of indicated ligands in cells expressing 53BP1 constructs. **h,** Total amount of indicated ligands (total fluorescence per focus) from same cells as in g. **i,** Proposed model of 53BP1 condensate foci as a bioreactor model. Larger condensate formation leads to increased NHEJ factors within an effective radius, increasing the effective concentration (C_eff_). Error bars in all panels, 95% CI. ns, not significant (p > 0.05); ***, p ≤ 0.001; ****, p ≤ 0.0001.

53BP1 is a molecular scaffold for the recruitment of additional machinery that participate in NHEJ^16, 21^. For example, 53BP1 recruits RIF-1 to DSBs (**Fig. 2e**) and their association has been shown to block inappropriate or alternative repair pathways^37–39^. Recruitment of the Shieldin complex at DSBs also contribute to repair activity through NHEJ ^40–42^. We first wanted to test whether the 53BP1 containing IDR1^RtoK^ mutations was capable of normal interactions with its binding partners. We measured the relative concentration of ligands localized to 53BP1 foci at break sites by co-immunofluorescence (**Fig. 2f**). We found similar levels of relative client concentration at 53BP1 foci for both wildtype and the IDR1^RtoK^ mutant (**Fig. 2g)** suggesting the mutant 53BP1, although deficient in self-assembly is nonetheless capable of normal recruitment of its key binding partners such as RIF-1 and Shieldin (Rev7), and likely other canonical ligands.

Having established normal adaptor function for the 53BP1 mutant, we next wondered whether reduced size of repair foci would cause a defect in the total recruitment of repair pathway ligand in proximity to a break site. Indeed, we found the total amount of ligand recruited to foci in the 53BP1-IDR1^RtoK^ mutant is significantly diminished (**Fig. 2h**). Taken together, these data suggest that 53BP1 compartments function to enforce the proximity of NHEJ factors to DNA breaks. If we consider the effective concentration of ligands within a distance of the break site (radius of wildtype condensates), 53BP1 condensates function as bioreactor of sorts that localizes NHEJ components above a threshold to promote DNA repair via NHEJ (**Fig. 2i**).

### 53BP1 condensation is required for efficient DNA repair

Our cellular observations suggest that robust 53BP1 condensation may be needed to recruit NHEJ factors within a regional volume of the break. To determine whether these mesoscale condensates are required for DSB repair, we quantified DSBs marked by γH2AX foci over time, following DNA damage induction by IR (**Fig. 3a**). We first compared 53BP1+/+ and 53BP1−/− cells synchronized in G1. Absence of 53BP1 profoundly delayed the resolution of γH2AX foci (**Fig. 3b, c and Extended Data Fig. 3a**), consistent with a central role for 53BP1 in DNA repair. We then compared stable cell lines expressing add-backs of wildtype 53BP1 versus the IDR1^RtoK^ mutant at the two-hour timepoint following damage (**Fig. 3d and Extended Data Fig. 3b**). Remarkably, while expression of wildtype 53BP1 suppressed persistent γH2AX foci similar to control cells, the IDR1^RtoK^ mutant was defective and showed similar number of unresolved γH2AX foci as the 53BP1 −/− deletion cell line. These data demonstrate that robust mesoscale condensation of 53BP1 is essential for efficient NHEJ DNA repair, and combined with our previous data suggest 53BP1 condensates function as bioreactors to enforce proximity of repair factors around DSBs.

**Fig 3.**
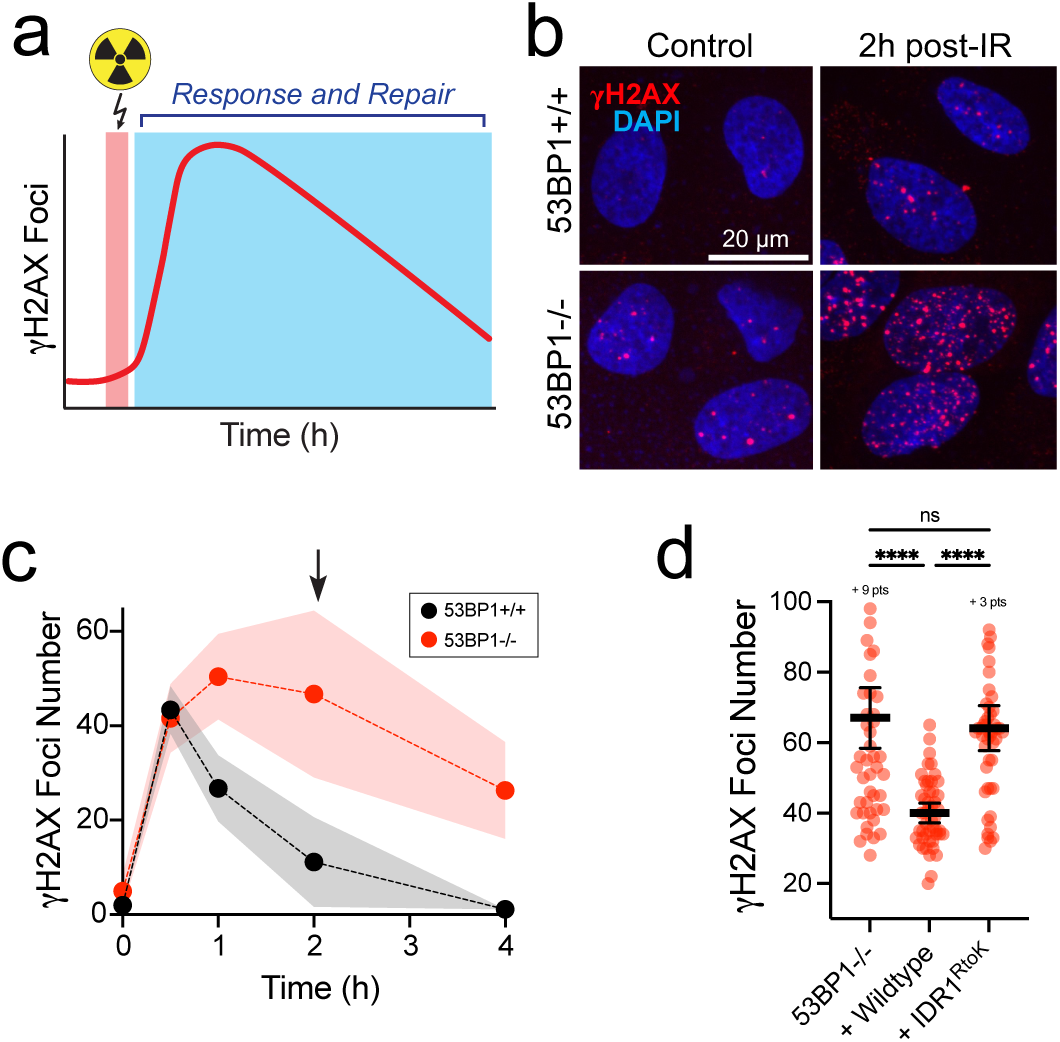
Condensation mutant of 53BP1 disrupts DSB resolution. **a**, Experiment schematic. Cells are subjected to 2Gy of ionizing radiation (IR), resulting in rapid accumulation of gH2AX foci that are resolved and disappear over time. **b,** Representative images of gH2AX immunofluorescence in nuclei with no IR exposure (control) and two hours post-IR from unperturbed and 53BP1−/− U2OS cells. **c,** Average number of gH2AX foci in immunofluorescence experiments as in b over time. Shaded area, 95% CI. Arrow indicates 2 h timepoint as showing largest dynamic range. **d,** Quantification of gH2AX foci number in 53BP1−/− and add-backs of full-length 53BP1 wildtype or R-to-K mutations in IDR1. Error bars, 95% CI. ns, not significant (p > 0.05); ****, p ≤ 0.0001.

### 53BP1 condensation insulates DSBs from aberrant end-processing and HDR in G1 phase

NHEJ and HDR repair pathways antagonize one another, and repair pathway choice depends on 53BP1 and key ligands such as RIF-1 and the Shieldin complex (**Fig. 2**). These factors bias repair toward NHEJ by blocking end processing^37, 38, 40^(**Fig. 4a**). While HDR is disfavored in G1 phase of the cell cycle due to the absence of replicated sister chromosomes^5^, loss of 53BP1 is associated with in increased HDR activity^43–45^.

**Fig 4.**
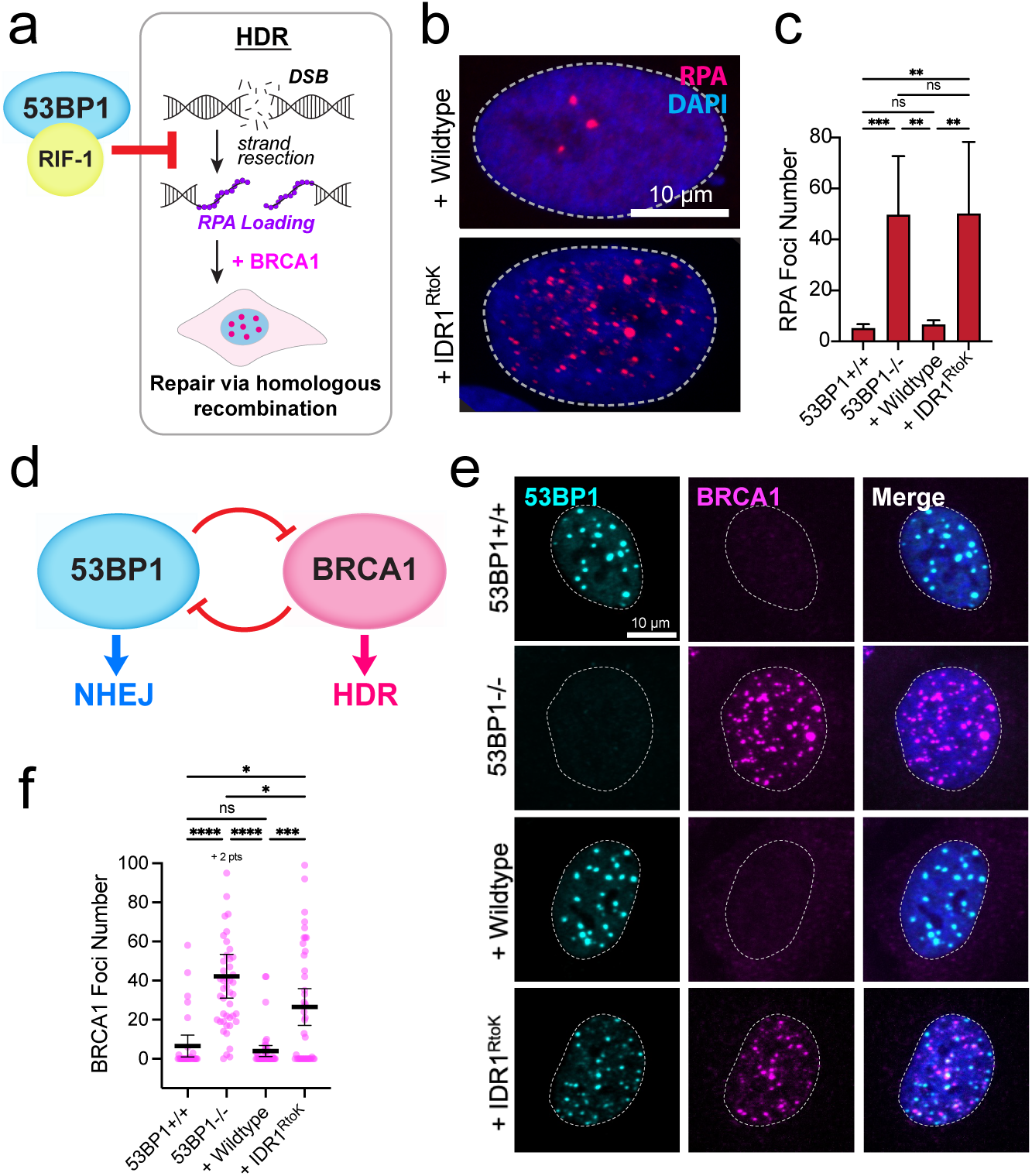
Diminished condensation for NHEJ leads to inappropriate HDR activation. **a**, Schematic. HDR required DNA strand resection, following a DSB. Single stranded DNA (ssDNA) is boundby RPA complex and BRCA1 complexes mediate repair via HDR. 53BP1 and ligands (e.g. RIF-1) block strand resection to prevent HDR. **b,** Representative IF images of RPA of nuclei expressing transduced constructs of 53BP1 wildtype or IDR1 mutant following DNA damage by IR. **c,** Quantification of average RPA foci number per cell in indicated cell lines. IDR1 mutant is unable to suppress RPA foci. **d,** Schematic of 53BP1 (NHEJ) and BRCA1 (HDR) antagonism. **e,** Representative IF images of 53BP1 and BRCA1 co-staining in the indicated cell lines. IDR1 mutant is unable to suppress BRCA1 foci, indicating HDR is initiated. **f,** Quantification of BRCA1 foci in cells as in e. Error bars in all panels, 95% CI. ns, not significant (p > 0.05); *, p ≤ 0.05, **, p ≤ 0.01; ***, p < 0.001; ****, p ≤ 0.0001.

It is unknown whether nanoscale recruitment of 53BP1 around DSBs – bound to chromatin – is sufficient to guide repair pathway selectivity or whether further mesoscale condensation of 53BP1 is instead needed to prevent end resection and block HDR. A key step in HDR initiation is DNA end resection stabilization of single stranded DNA by the RPA complex^46–48^ (**Fig. 4a**). To assess the presence of resected DNA, we measured the number of RPA foci in cells following IR in G1 (**Fig. 4b, c and Extended Data Fig. 4**). 53BP1+/+ cells have few RPA foci. Loss of 53BP1 results in numerous RPA foci, indicating resection and HDR initiation (**Fig. 4c**). Stable addition of wildtype 53BP1 into 53BP1−/− cells suppressed RPA foci formation similar to unperturbed cells. However, 53BP1 harboring IDR^RtoK^ mutations is unable to suppress RPA foci (**Fig. 4b, c**). These data demonstrate that 53BP1 condensate formation at DSBs is required to block end-resection and bias repair pathways choice to NHEJ.

53BP1 and BRCA1 antagonize each other^6, 11, 43, 49^ (**Fig. 4d**) and their associated machinery are expressed at similar levels in all cell cycle phases^39^. Further, knockdown of 53BP1 results in foci aberrant BRCA1 foci and vice versa^43^. This is consistent with a model in which NHEJ and HDR are in competition across the cell cycle. To test whether 53BP1 condensate formation is required to suppress BRCA1 activity, we measured the presence of BRCA1 foci in cell lines synchronized in G1. Unperturbed 53BP1+/+ cells had little or no visible BRCA1 foci in G1 (**Fig. 4e, f**). Consistent with previous observations, loss of 53BP1 results in increased HDR and formation of many large BRCA1 foci^43^, even in G1 (**Fig. 4e, f, g**). Reintroduction of wildtype 53BP1 effectively suppresses BRCA1 foci similar to the unperturbed control (**Fig. 4f**). The presence of the IDR1^RtoK^ mutant of 53BP1 is not able to effectively suppress BRCA1 foci formation (**Fig. 4e, f**).

Taken together, our work demonstrates nanoscale clustering of 53BP1 is not sufficient to block HDR. Instead, robust condensate formation by 53BP1 is essential for repair pathway choice by blocking end resection and insulating against HDR foci formation around DSBs. Future studies will be required to determine whether this insulation activity occurs through functional blocking (ie RIF-1 antagonizes HDR activities) or physical insulation, such as blocking diffusion of HDR components within proximity of DSBs.

## DISCUSSION

In this study, we identify the mechanism by which repair foci for end-joining are assembled through biochemical reconstitution and cell culture experiments, revealing an essential role for repair protein condensation in control of genomic integrity. To our knowledge, this is the first study to demonstrate a direct link between condensation and efficiency of the repair process in cells. We identified the minimal domain necessary and sufficient for 53BP1 biomolecular condensation and determined the amino acids that govern its phase separation. The minimal region includes the oligomerization domain and C-terminal IDR1 of 53BP1. The assembly of a diverse array of condensates is driven by proteins whose domain architecture resembles 53BP1, containing an OD and IDR, and ligand binding sequences^24^. Oligomerization activity of 53BP1 has previously been implicated in several of its functions including DNA repair, class switch recombination, and chromatin maintenance^50–52^. We found the IDR1 sequence was enriched in R residues, interactions of which have been shown in multiple studies to play central roles in protein condensation^30, 31, 53, 54^. Mutation of a number of these R residues have also been associated with cancer development^55^. R-to-K mutations in IDR1 abolished phase separation in vitro and provided a tool to dissect the role of 53BP1 condensation in cells. Full-length 53BP1, containing IDR1 R-to-K mutations, formed smaller foci after DNA damage and caused defects in DNA repair. The requirement of condensation for repair efficiency was revealed by delayed kinetics in resolution of breaksites, aberrant appearance of RPA on inappropriately resected DNA, and inappropriate formation of HDR foci containing BRCA1 during G1 phase of the cell cycle.

Protein self-assembly and biomolecular condensate formation is a ubiquitous form of subcellular compartmentalization^28, 56, 57^. Over the past decade, protein condensates have been implicated in numerous nuclear processes including ribosomal biogenesis within nucleolar bodies, chromatin compaction through condensation HP1α, transcription at super-enhancer assemblies governed by Mediator, telomere clustering, and more recently in maintenance of genomic integrity through tumor suppressors and DNA repair enzymes^19, 35, 58–66^. In each case, these protein ensembles support selective partitioning of machinery involved in an assigned nuclear process. In essence, biomolecular condensates act to increase the effective concentration of enzymes to support a specific function. Similarly, synthetic condensate systems have been developed as compartments that enhance local concentrations of effectors^67–70^. Our data support this view that 53BP1 ensembles compartmentalize their associated machinery at break sites for NHEJ. The IDR1^RtoK^ mutant is unable to form condensates in vitro, and full length 53BP1 containing IDR1^RtoK^ mutation forms smaller condensates in cells without affecting binding partner interactions. Reduced sizes of 53BP1 foci were strongly linked to reduced functionality, supporting the idea that an effective concentration of recruitment is needed to efficiently drive NHEJ, consistent with a bioreactor model for condensate function.

Our data also suggest that 53BP1 condensates simultaneously function as insulators, preventing other competing pathways, either physically or functionally from accessing break sites. Condensates including Balbiani bodies, stress granules, p-bodies, as well as engineered condensate systems have been shown to selectively sequester proteins and nucleic acids, thus preventing access to other interactors^71–80^. Our findings suggest that 53BP1 condensates prevent end-resection machinery and BRCA1 foci from accessing repair sites. We show 53BP1 condensation in G1 phase of the cell cycle is needed to block RPA accumulation (a marker for resection) and assembly of foci containing BRCA1. Thus, insulator function of 53BP1 condensates prevents hallmarks of the HDR response and represents a major basis for pathway competition in G1.

Collectively, this study sheds light on an elusive function of 53BP1. While its C-terminus was found to phase separate in vitro^19^, the mechanism by which 53BP1 self-assembles and the specific role that 53BP1 condensate formation plays in DNA damage response has been unclear. Although 53BP1 does not participate in physically ligating broken ends^81–83^ it plays a key role in end protection to enable NHEJ action. 53BP1 self-assembly and condensations serves to create a mesoscale compartment to recruit binding partners while additionally insulating against BRCA1 foci and HDR repair machinery that have the potential for mutagenic repair in G1. This work contributes to an emerging paradigm that to safeguard genome integrity, factors must selectively partition within membraneless compartments.

## AUTHOR CONTRIBUTIONS

MVG, RAG, MCG, and TZ conceived of the project and experiments. MVG and AD performed the experiments. WW and MW provided experimental support, performed cell sorting, and cloning. MVG wrote the manuscript. MVG, RAG, and MCG edited the manuscript.

## ACKNOWLEDGEMENTS.

We thank and Dr. Haoyang Jiang for advice and technical support. We thank the Penn Center for Genomic Integrity for feedback on this project. We also thank Dr. Andrea Stout and the Penn CDB Microscopy Core for training, imaging, and support. This study was supported by a National Institute of Biomedical Imaging and Bioengineering grant, (EB028320) and by the NSF through the University of Pennsylvania Materials Research Science and Engineering Center (MRSEC) (DMR-2309043) to MCG. This work was also supported by an American Cancer Society Postdoctoral Fellowship (PF-21-186-01-DMC) and a K99/R00 Award from the National Institute of General Medical Sciences (K99GM151473) to MVG. Additionally, this work was supported by a grant from the National Cancer Institute (R01 CA174904) to RAG.

## METHODS

### Cloning

All plasmids were constructed using InFusion cloning (Takara Bio; Mountain View, CA) and verified by Plasmid-EZ and/or Sanger sequencing (Azenta Life Sciences; Burlington, MA). Sequences for protein expression and purification were codon optimized for bacterial expression and ordered as gene fragments (Twist Bioscience; San Francisco, CA). Sequences were amplified by PCR and cloned into pET or pMALc2 vectors. Lentiviral donor plasmids were made in the pLJM1 vector from PCR amplifications of the 53BP1 N-terminus from a pMX-53BP1 plasmid (gift from the Greenberg Lab) and 53BP1 C-terminus from the same plasmid or from synthesized gene fragments (Twist Bioscience).

### Cell culture

U2OS human osteosarcoma and HEK293T cell lines were cultured in Eagle’s Minimal Essential Medium (EMEM; Quality Biological) supplemented with Gibco 10% Fetal Bovine Serum (Thermo Fisher Scientific; Waltham, MA), 2 mM L-glutamine (Gibco), and 5 µg/mL Plasmocin (InvivoGen; San Diego, CA). All cell cultures were maintained at 37°C in a humidified atmosphere with 5% CO_2_. Cells were split in a 1:5 ratio every 3 days and were limited to 25 passages. Cell lines were negative for mycoplasma infection, and all experiments were performed with confirmed viability >95% by Trypan Blue staining (Thermo Fisher Scientific). The 53BP1−/− U2OS cell line was a gift from the Greenberg lab, in which both copies of 53BP1 were previously deleted via CRISPR.

To generate stable cell lines, lentiviral donor plasmid encoding a GFP-53BP1 variant (wildtype or IDR1 mutant) were co-transfected with pMD2.G (VSV-G envelope plasmid; a gift from Didier Trono – Addgene plasmid # 12259; http://n2t.net/addgene:12259; RRID:Addgene_12259) and psPAX2 (lentivirus packaging plasmid; a gift also from Didier Trono – Addgene plasmid # 12260; http://n2t.net/addgene:12260; RRID:Addgene_12260) into HEK293T cells at approximately 70% confluency in a 10 cm cell culture dish (Corning). Briefly, 10 µg of the lentiviral donor plasmid, 2.5 µg pMD2.G, and 7.5 µg psPAX2 were incubated in 250 µL of Opti-MEM (Thermo Fisher Scientific) for 5 min at ambient temperature (∼ 22°C). Simultaneously, 60 µL Lipofecatmine-2000 (Thermo Fisher Scientific) was incubated in a separate 250 µL Opti-MEM for 5 min at ambient temperature.

Mixtures were then combined and incubated for 15 min at ambient temperature. Mixtures were then applied dropwise to HEK293T cells and incubated for approximately 16 hrs at 37°C in a humidified atmosphere with 5% CO_2_. EMEM media was changed and transfected cell cultures were incubated for an additional 36 hrs. Media containing virus was collected and concentrated by centrifugation in Amicon centrifugal filters with a 200 kDa cutoff (MilliporeSigma; Burlington, MA), centrifuged for 5-10 min at 1,500 x g in a benchtop swinging bucket centrifuge (Applied Biological Materials; Richmond, BC, Canada).

Concentrated virus was used immediately to transduce 53BP1−/− U2OS cells seeded in a 24-well cell culture dish (Greiner Bio-One). Cell culture media was replaced with virus containing media supplemented with 10 µg/mL Polybrene (MilliporeSigma). Transductions were incubated at 37°C in a humidified atmosphere with 5% CO_2_ for approximately 16 hrs, after which media containing virus was replaced with fresh EMEM and incubated for 2-3 days. Cells expressing 53BP1 were isolated via cell sorting with an Aria C cytometer (BD Biosciences). The IDR1 mutant harbors 13 R-to-K mutations, avoiding mutation of arginines located in the Gly/Arg-rich (GAR) motif as they may be involved in 53BP1 targeting^84, 85^.

### Immunofluorescence

U2OS cell lines were seeded in an 8-well cell culture imaging slide (CellVis; Mountain View, CA) and incubated overnight at 37°C in a humidified atmosphere with 5% CO_2_. Cells were then synchronized in late G1 phase with media supplemented with 10 µM Palbociclib (MedChemExpress; Monmouth Junction, NJ) for 24 hrs. The drug was then washed out and replaced with fresh media and incubated for 40-60 min. DNA damage was induced by exposing cells to 2 Gy ionizing radiation (IR) in a Cs-137 Gammacell Irratior (Nordion; Ottowa, Canada). Cells were then incubated at 37°C in a humidified atmosphere with 5% CO_2_ until specified timepoints. Cells were then washed three times, for 5 min each, with phosphate buffered saline (PBS) and fixed in fixation buffer (3% formaldehyde and 2% sucrose in PBS) for 10 min at 37°C. Cells were washed three times, for 5 min each, in PBS to remove fixation buffer and permeabilized with 0.5% Triton X-100 in PBS for 10 min at ambient temperature. Cells were washed three times in PBS and blocked for 1 hr at ambient temperature in blocking buffer (1 mg/mL BSA, 3% FBS, 0.1% Triton X-100, pH adjusted to 7.5). Cells were then incubated with blocking buffer containing primary antibodies (**Supplementary Table 1**) and 0.02% NaN_3_ in a humidified chamber overnight at 4°C. Following primary antibody incubation, cells were washed three times, for 5 min each, in PBS containing 0.5% Tween-20 at ambient temperature. They were then incubated in blocking buffer with secondary antibody (**Supplementary Table 2**) for 1 hr at ambient temperature. Following secondary antibody incubation, cells were washed three times, for 5 min each, in PBS. Chromatin was stained with DAPI (1 µg/mL in PBS) for 5 min and washed twice in PBS, 5 min each. Cells were imaged immediately.

Fluorescence microscopy of stained cells was performed at ambient temperature on an Olympus IX83 inverted microscope (Olympus Life Science; Tokyo, Japan) equipped with a CrestOptics X-Light V3 spinning disk (CrestOptics; Rome, Italy), multiple excitation laser launches (405, 445, 488, 520, 555, and 630 nm), and two pco.edge 4.2 bi sCMOS cameras (Excelitas; Pittsburgh, PA) under the control of VisiView software (Visitron; Puchheim, Germany). The microscope stage is enclosed in an Okolab incubation chamber (Okolab; Pozzuoli, Italy). Samples were illuminated and images collected through a 60x/1.3 NA silicone-immersion objective. Black and white values for images shown were manually adjusted in Fig 2 due to differences in expression or differences in antibody or DAPI staining.

### Protein purification

Recombinant proteins of wildtype or mutants of all R residues in IDR1 to K were purified from bacteria. In all cases, plasmids for protein expression were transformed into Rosetta2 bacterial cells harboring the pRARE plasmid and grown on Luria Broth (LB) plates supplemented with chloramphenicol (to maintain the pRARE plasmid) and either carbenicillin or kanamycin (to maintain protein expression vector) at 37°C overnight. A single colony was then grown in 2 mL LB supplemented with the appropriate drugs to saturation. This was then transferred to 100 mL fresh media and grown to saturation. From this, 25 mL was transferred to each liter of fresh media (2-4 L per protein expressed) and grown at 37°C to OD_600_ or 0.6 to 0.8. Protein expression was induced with 0.5 mM Isopropyl β-D-1-thiogalactopyranoside (IPTG) overnight at 16°C. Bacterial pellets were collected by centrifugation at 4,500 x g in a Sorvall RC6+ centrifuge (Thermo Fisher Scientific) and stored at –80°C.

To purify expressed polypeptides, bacterial pellets were thawed and resuspended in lysis buffer (50 mM Tris-HCl, pH 7.5, 1 M NaCl, 20 mM imidazole, 1 mM β-mercaptoethanol) and complete EDTA-free protease inhibitor cocktail (Roche; Mannheim, Germany). Cells were lysed with 1-2 minutes of sonication at 50% amplitude using a Branson Sonifier. Lysates were clarified by centrifugation at 18,000 x g for 25 min in a Sorvall RC6+ centrifuge using a F21S-8×50y rotor (Thermo Fisher Scientific). Supernatants were incubated with 0.5 mL of Ni–NTA beads (Thermo Fisher Scientific) while rotating at 4°C for 1 hour. Beads were then washed three times with 10 mL in wash buffer (20 mM Tris-HCl, pH 7.5, 0.5 M NaCl, 20 mM imidazole, and 1 mM β-mercaptoethanol). Proteins were eluted in elution buffer containing 20 mM Tris-HCl, pH 7.5, 0.5 M NaCl, 20 mM imidazole, and 1 mM β-mercaptoethanol. For polypeptides fused to an N-terminal MBP tag, Ni-NTA elutions were diluted 5-10 fold in buffer lacking imidazole and incubated with 0.5 mL amylose beads while rotating at 4°C for 1 hour. Beads were then washed three times with 10 mL in wash buffer (20 mM Tris-HCl, pH 7.5, 0.5 M NaCl, and 1 mM β-mercaptoethanol) and then eluted with the same buffer supplemented with 10 mM maltose.

Eluted proteins were diluted to approximately 3 mg/mL in buffer lacking imidazole or maltose and dialyzed overnight into storage buffer (20 mM Tris-HCl, pH 7.5, 500 mM NaCl, 1 mM DTT) using Slide-A-Lyzer membrane cassette (Thermo Fisher Scientific) with a 20 kDa size cutoff at 4°C. Proteins were concentrated by centrifugation in 4 mL of Amicon filter concentrators. (Millipore Sigma; Burlington, MA). TCEP (1 mM) was added prior to snap freezing and storage at –80°C.

### In vitro condensate assembly

To test protein droplet formation in vitro, purified polypeptides were thawed at 50°C and diluted buffer to a final salt concentration of 150 mM NaCl and 20 mM Tris-HCl pH, 7.5. For MBP tagged constructs, the MBP tag was cleaved off the polypeptide of interest with TEV protease at a 20 to 1 molar ratio of MBP tagged protein to TEV protease for 30 min. These mixtures were applied to custom fabricated acrylic gasket chambers adhered to glass cover slips at ambient temperature. Protein condensate formation was observed within 30 min by microscopy. Imaging of in vitro protein droplet formation was performed at ambient temperature on an Olympus IX81 inverted confocal microscope (Olympus Life Science; Tokyo, Japan) equipped with a Yokogawa CSU-X1 spinning disk and an iXon3 EMCCD camera (Andor; Belfast, UK). Multidimensional image acquisition was controlled by MetaMorph software (Molecular Devices; San Jose, CA). Samples were illuminated using a brightfield lamp and imaged through a 100x/1.4 NA oil-immersion objective.

### Analysis

Sliding window analysis in Fig. 1e was performed with a custom written script in Python. Data are binned in 50 residue blocks with a 5 amino acid sliding window and quantify the percentage of arginines. For image analysis, quantification of foci number and volume was performed via custom macros in ImageJ. Nuclei were segmented using the DAPI channel and masks for each nucleus were generated by automated thresholding. Masks were then applied to MAX intensity projections of fluorescence images and individual nuclei were cropped. Masks for foci were then generated from each nucleus image to quantify foci number and area (µm^2^). Area was converted to volume (µm^3^) by cubing the square root of the foci area.

### Statistics and Reproducibility

Experiments were generally performed 2-3 times and were reproducible. All statistical analyses were performed in GraphPad Prism 9. To test the significance of two categories, an unpaired two-tailed t-test was used. To test significance of more than two categories, a one-way ANOVA was used. Results in graphs are shown as mean values and error bars are 95% CI. In all cases: ns, not significant; P > 0.05, * P ≤ 0.05; ** P ≤ 0.01; *** P ≤ 0.001; **** P ≤ 0.0001.

### Data Availability

All data used in this study are available from the corresponding author upon reasonable request. Macros and scripts used for sliding window analysis or to automate image analysis will be made available on Github.

---

**Extended Data Fig. 1.**
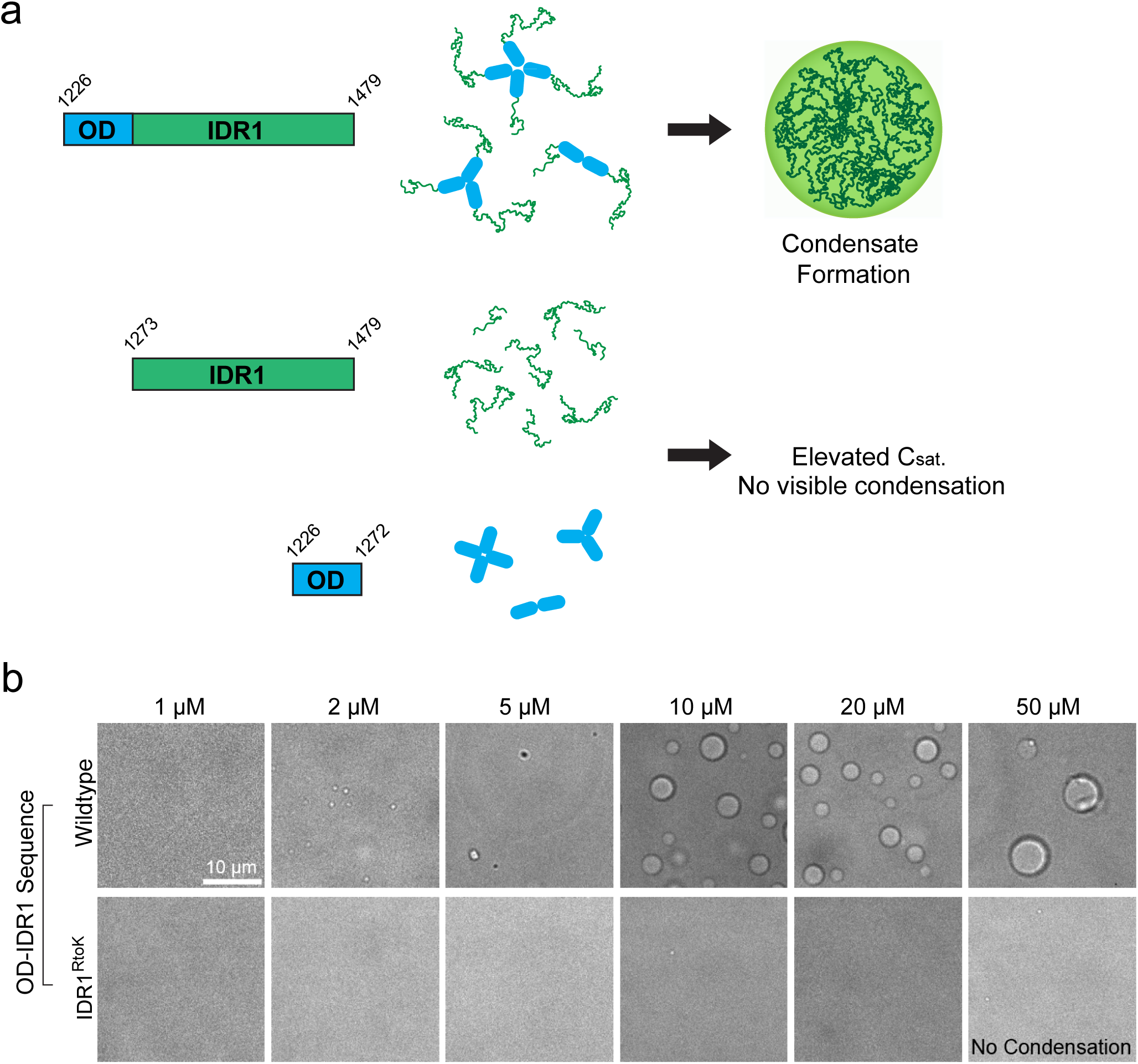
**a**) Diagrams of 53BP1 OD-IDR1 (1226-1479), IDR1 (1273-1479), or OD (1226-1272) sub-fragments. OD-IDR1 is capable of condensation. However, IDR1 alone does not condense at the tested concentrations, likely due to elevated C_sat_. OD alone may cluster, but does not produce visible condensed structures. **b)** Comparison of in vitro condensation of wildtype OD-IDR1 and OD-IDR1 RtoK mutant in which arginines (R) in the IDR1 sequence are mutated to lysine (K). Lysines maintain residue charge but break arginine interactions.

**Extended Data Fig. 2.**
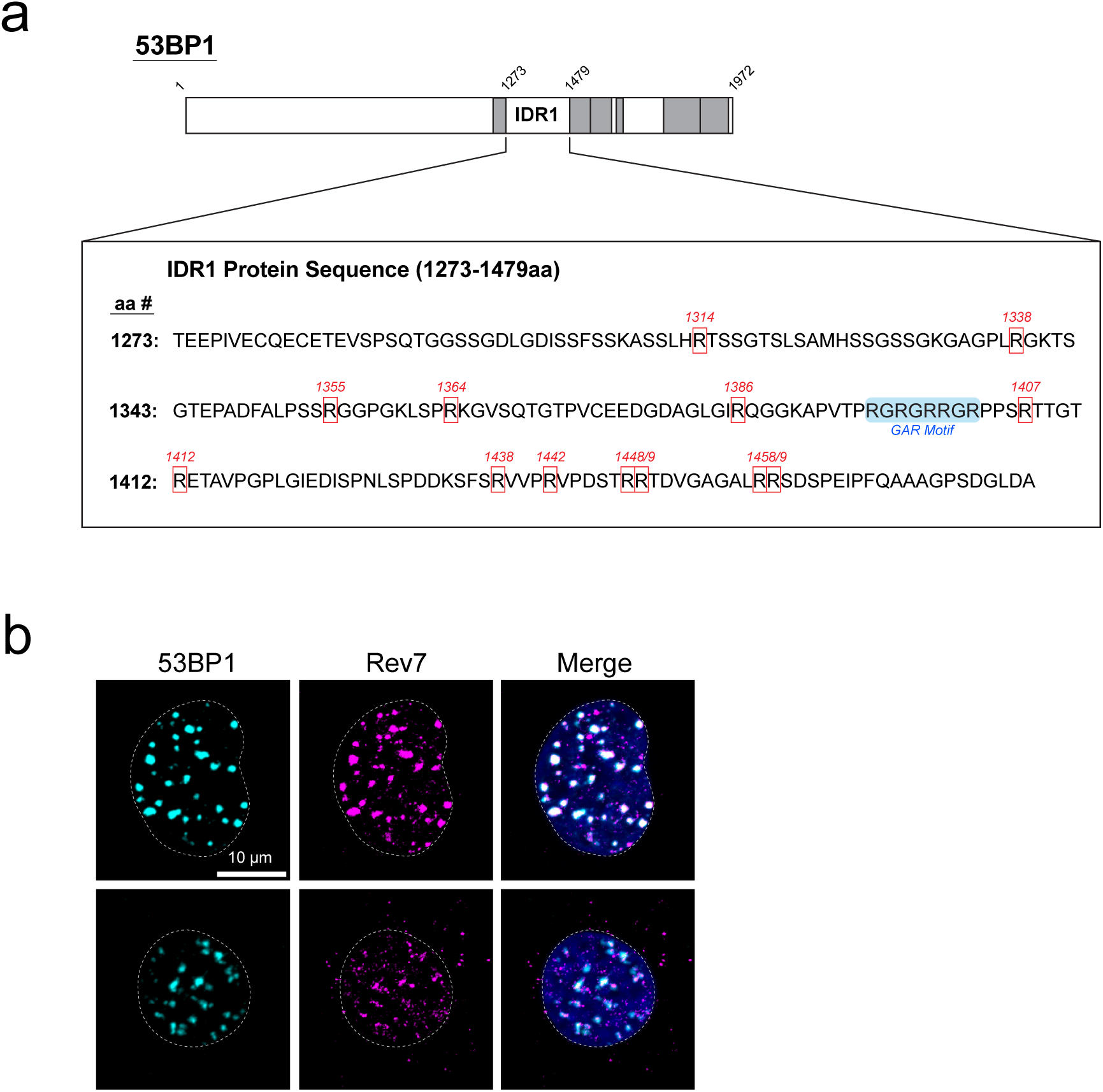
**a**) Diagram of 53BP1 and amino acid sequence of its C-terminal IDR1 (1203-1479). Arginines mutated to lysines (R-to-K) in 53BP1 constructs transduced into 53BP1−/− cells are boxed in red. Arginines located within the Gly/Arg-Rich (GAR) motif are highlighted in blue and were not mutated. **b)** Representative immofluorescence images of Rev7 (component of the Shieldin Complex) and 53BP1 in stable cell lines expressing wildtype or mutant 53BP1.

**Extended Data Fig. 3.**
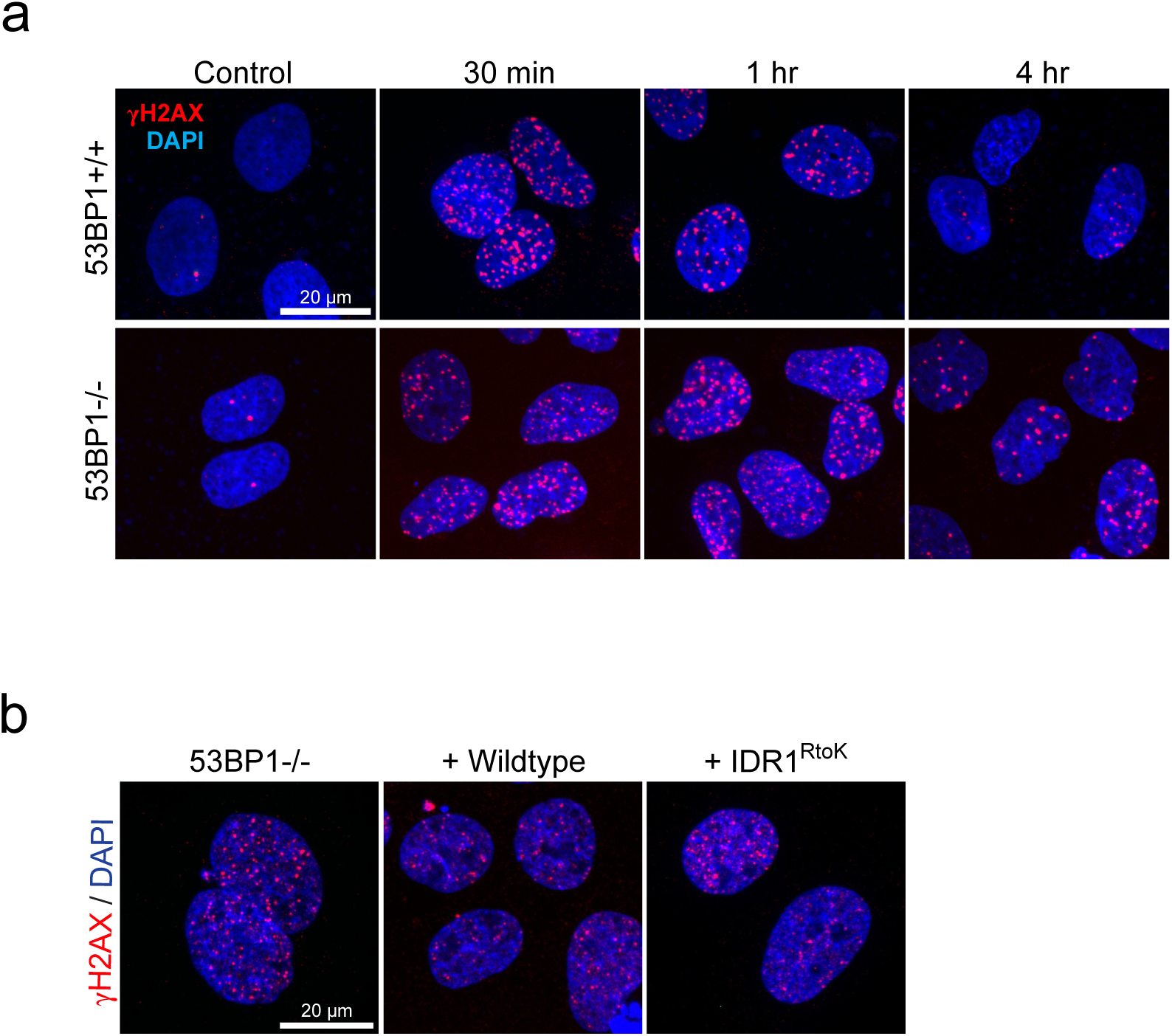
**a**) Representative immunofluorescence images of gH2AX foci (marking DNA DSBs) in nuclei over time following 2 Gy IR. Loss of 53BP1 results in reduced resolution of gH2AX foci compared to 53BP1+/+. **b)** Representative immunofluorescence images of gH2AX foci 2 hrs following 2 Gy IR in 53BP1−/− cells and add backs of wildtype and mutant (IDR1^RtoK^) 53BP1. Wildtype add back resolves foci while mutant is similar to 53BP1−/−.

**Extended Data Fig. 4.**
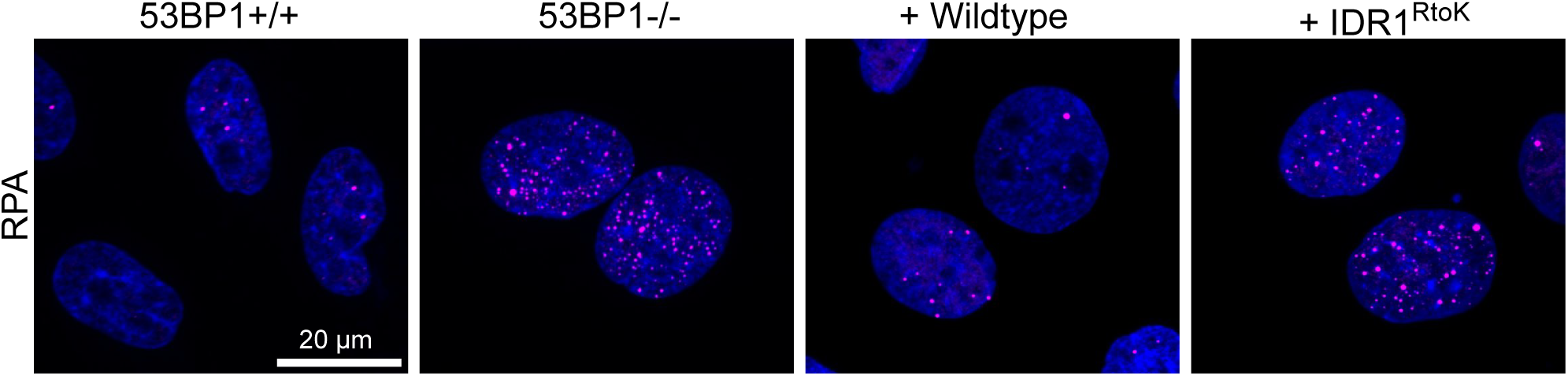
Representative immunofluoresence images of RPA foci in nuclei of indicated cell lines, synchronized in G1 and exposed to 2 Gy IR. 53BP1+/+ cells suppress RPA foci while 53BP1−/− is unable to suppress RPA foci formation. Add back of wildtype 53BP1 restores suppression of RPA while the condensation mutant, IDR1^RtoK^, is unable to do so.

