## Supplementary Materials PDF for "53BP1 condensates function as bioreactors for NHEJ directed DNA repair and insulators to determine pathway choice"

**53BP1 condensates function as bioreactors and insulators at DNA breaks to promote non-homologous end joining.**

**This PDF file includes:**

Supplementary Table 1

Supplementary Table 2

**Supplementary Table 1.**

| Primary Antibody | Concentration for IF | Source |
| --- | --- | --- |
| Rb α γH2AX | 1:1000 | Active Motif |
| Ms α 53BP1 | 1:1000 | Novus |
| Rb α 53BP1 | 1:1000 | Novus |
| Rb α RIF-1 | 1:1000 | Novus |
| Rb α Rev7 | 1:200 | Abcam |
| Ms α RPA2 | 1:500 | MilliporeSigma |
| Rb α BRCA1 | 1:1000 | SantaCruz |

**Supplementary Table 2.**

| Secondary Antibody | Concentration for IF | Source |
| --- | --- | --- |
| Gt α Rb AlexaFluor 488 | 1:2000 | Thermo Fisher Scientific |
| Gt α Ms AlexaFluor 488 | 1:2000 | Thermo Fisher Scientific |
| Gt α Rb AlexaFluor 568 | 1:2000 | Thermo Fisher Scientific |
| Gt α Ms AlexaFluor 568 | 1:2000 | Thermo Fisher Scientific |
| Gt α Ms AlexaFluor 640 | 1:2000 | Thermo Fisher Scientific |

**Abbreviations Used.**

- **Rb:** rabbit
- **Ms:** mouse
- **Gt:** goat
- **α:** anti
